# Imputation of posterior linkage probability relations reveals a significant influence of structural 3D constraints on linkage disequilibrium

**DOI:** 10.1101/255315

**Authors:** Susanne Gerber, David Fournier, Charlotte Hewel, Illia Horenko

**Affiliations:** Center for Computational Sciences in Mainz (CSM) and Institute for Developmental Biology and Neurobiology at the Faculty of Biology, Johannes-Gutenberg University of Mainz, Staudinger Weg 9, 55128 Mainz, Germany; bInstitute of Computational Science, Faculty of Informatics Università della Svizzera Italiana, Via G. Buffi 13, 6900 Lugano, Switzerland

**Author notes:** The authors declare that they have no competing interests.

**Keywords:** Relation measure, Genome Wide Association Studies, Multiscale processesi, Information criteria, Numerical methods

## Abstract

Genetic association studies have become increasingly important in unraveling the genetics of diseases or complex traits. Despite their value for modern genetics, conflicting conclusions often arise through the difficulty of confirming and replicating experimental results. We argue that this problem is largely based on the application of statistical relation measures that are not appropriate for genomic data analysis and demonstrate that the standard measures used for Genome-wide association studies or genomics linkage analysis bear a statistic bias. This may come from the violation of underlying assumptions (such as independence or stationarity) as well as from other conceptual limitations in the measures or relations, such as missing invariance with respect to coding or the inability to reflect latent factors. Attempts to introduce unbiased relation measures that avoid these limitations are usually computationally expensive and do not scale for large data sizes being typical for genomics applications.

To tackle these problems, we propose a straightforwardly computable relation measure called Linkage Probability (LP). This measure provides the posterior probability of a relation between two categorical data sets and considers potential biases from latent variables. We compare several aspects of popular relation measures through an illustrative example and human genomics data. We demonstrate that the application of **LP** to the analysis of Single Nucleotide Polymorphisms (SNP) reveals latent 3D steric effects within 1D SNP data, that approximate to chromatin loops captured by high resolution Hi-C maps.

---

One of the central goals of personalized medicine is to develop methods of diagnostics to establish robust links between patient-specific information and indicators relevant to therapy (disease risk, probabilities of complications from therapies, a susceptibility to certain drugs, etc.) (1). An important problem in this context is the evaluation of the patient’s genetic background. Genome Wide Association Studies (GWAS) are a powerful tool that permits linking traits or diseases to underlying genetics (e.g. genes, Quantitative Trait Locations (QTL) or common point mutation (Single Nucleotide Polymorphisms, or SNP) (2–4). Another approach to analyzing the principles of genetic variants is the computation of Linkage Disequilibrium (LD), defined as the nonrandom association of alleles (variant forms of a given gene) at different gene loci in a given population. Especially through the dramatic increase of the number of genomes that have become available over the past decade, LD has become a key approach in the analysis of relations between diverse SNP to make new genetic discoveries related to the pathogenesis of diseases and disorders (5–8). LD can be used either to establish pairwise associations between SNP to specific diseases, or, to infer genetic, evolutionary and chronological relationships between SNP of different species and populations

Generally speaking, this problem can be seen as the computation of relation for a pair of variables, *X* and *Y*, where the value of *Y* is obtained from *m* categories *y* = {*y*(1),*y*(2),…,*y*(*m*)} and the value of *X* derived from *n* categories *x* = {*x*(1),*x*(2),…,*x*(*n*)}. Each set of the SNP datasets {*X*^1^,…,*X*^*T*^} and {*Y*^1^,…,*Y*^*T*^} represent allele states for two particular SNP that were collected from *T* subjects ( with *m* = 3 genotype classes and *x* = *y* = {*’homozygous major’, ‘heterozygous’ and’homozygous minor’*}).

In this article, we will demonstrate that common measures currently used to detect pairwise relationships between SNP are either biased or have at least a strong potential to introduce errors due to their non-robust nature. Since the discussion of the various sources of errors in the currently available measures are not only essential to this article, but may also expose a potential bias in the results from hundreds of studies based on them, we will highlight the major sources of such biases here, as well as in the Supplement, in greater details.

## Significance Statement

Using a very simple example we show that non of the popular measures used in genomics and in Genome-Wide Association Studies (GWAS) are unbiased. We propose a novel computationally-scalable measure for extracting the linkage patterns from categorical data and illustrate its benefits with a human genome data application. Compared to the standard measures (e.g., cross-correlation and linkage disequilibrium), this measure is designed to explicitly account for an eventual implicit impact from latent variables - and, in contrast to the measures based on the p-value concept, it provides a direct estimate of the posterior probability of a relation conditioned on the given data. Provided results exemplify that this new measure helps to capture the latent 3D epigenetic/steric effects based on the 1D SNP data only - by revealing the implicit/latent proximity of SNP in 3D space. We also show that from the conceptual view point this new measure - being based on the exact law of the total probability – is coding-invariant. We believe that this measure has a chance to become one of the main workhorses in GWAS and genomic studies.

**The Linkage Disequilibrium measure, LD** between two categories *i* and *j* for variables *Y*^*t*^ and *X*^*t*^ has been used since the first GWAS studies and remains one of the most popular methods of capturing correlations between pairs of SNP associated to a condition. This approach measures the nonrandom association of alleles at different loci (9). The level of linkage disequilibrium between *Y*^*t*^ and *X*^*t*^ can be quantified by the coefficient *D*_*ji*_ = ℙ [*X*^*t*^ = *x*(*j*) and *Y*^*t*^ = *y*(*i*)] – ℙ [*X*^*t*^ = *x*(*j*)] ℙ [*Y*^*t*^ = *y*(*i*)] (3, 10), which measures the difference between the frequency of the co-occurence of two SNP within a population and the assumption that the two loci mutate independently one from another. The calculation of this coefficient *D*_*ji*_, however, reflects an assumption that for each pair of categories *y*(*i*) and *x*(*j*) for *m* > 2 only two models: total dependence and total independence. Therefore, LD does not allow to study all possible dependence and independence combinations in all of the categories simultaneously. A demonstration that this introduces bias into classical LD measures can be seen in another case as well, for example when a variable *Y* comprises more than two categories (see section 2 in the Supplemental Information for "Material and Methods"). Classical LD and variations thereof (see (3, 11, 12) for comprehensive reviews) suffer from additional weak points: they are strongly influenced by allele frequencies, which are biased if populations are stratified, or influenced by other confounding factors; additionally, as mentioned above, they are restricted to pairwise comparisons and thus not cannot cope with marker data from multiple loci. Despite these issues, LD has served as the basis of a number of recent high-profile studies and are implemented in state-of the art software packages such as PLINK (13), INTERSNP (14) and Haploview (15).

**Latent factors distort conclusions regarding SNP relationships** For the most part, the measures currently used in practical applications for categorical variables (such as A, G, C, T) can be subdivided into three general classes:

- a correlation/covariance measure class based on Euclidean distances (e.g., Euclidean measure, Mahalanobis distance, cross-correlation measures, linear regression measures as implemented in INTERSNP, Granger causality measure, etc.) (14, 16–18);
- a measure family based on the processing of contingency tables (e.g., χ^2^-test, Fischer’s exact test and Fischer’s relation measure, Cramer’s V-measure, Bernoulli or Markov measures, etc.) (3, 6, 19)
- measures from information theory (e.g., Hamming distance, Jaccard index, Levenshtein distance (20, 21)).

Despite their diversity, all these measures exhibit a common weakness: the evaluations obtained from these standard statistical measures. In assigning a measure of of significance/insignificance of certain relationships and drawing conclusions about their strength or weakness, conclusions can be spoiled by latent factors (22). This means that the specific values *t* obtained when considering variables *Y*^*t*^ and *X*^*t*^ might be affected by other variables *U*^*t*^ that are not explicitly presented as data, where *t* = 1,…,*T* is an index of realization and *T* is the sample size. While the nature and type of influence that such latent variables *U*^*t*^ exert may be unknown, they can directly affect potential associations between *Y^*t*^* and *X*^*t*^, and changes in *U*^*t*^ may alter them, introducing a *t*-dependence of *Y* and *X*. Explicitly incorporating such effects into computations of standard measures might generate ill-posedness and overfitting problems (23, 24). In the example of an analysis of genomics data presented below, *U*^*t*^ may be some other SNP or genes that affect the relationship between two particular SNP *X*^*t*^ and *Y*^*t*^ as well as other factors, such as sex, the 3D geometrical structure of chromatin, the methylation status of a gene, a dependence on time or environmental factors that are specific to an individual t. Even though no one denies that such factors have strong effects on a phenotype or the relationship between specific variables that have been observed, they are systematically neglected, and this leads to a similar bias in the results of an analysis.

**The explicit incorporation of strong a priori assumptions on the underlying derivation of data** is an additional problem of standard measures that lead to biased results if the assumptions are not fulfilled. To provide an example, correlation measures explicitly rely on the Euclidean distance, which is not a proper metric for categorical variables and can vary with respect to a renaming or renumbering of categories, as illustrated below. A prerequisite for applying this measure to categorical data is first that the data must undergo a transformation from categorical into Euclidean space, for example by assigning some real number to every category. However, as the number possible transformations is theoretically infinite, the results from applying this transformation will be neither unique nor robust. As demonstrated in an example below (see Table 1), this can lead to a range of results ranging from strong negative correlation and independence to a strong positive correlation; the result is entirely determined by the user’s choice of transformation. We elaborate on and demonstrate this conceptual problem of standard measures on an illustrative toy example presented below. For the same reason, this problem extends further, to applications of methods such as linear regression, Principal Component Analysis (PCA) and parametric methods of Gaussian statistics (all of which are based on the Euclidean distance), to issues in categorical data analysis and reductionism in genomics. This means that such widely-used standard approaches may produce biased and non-unique results when applied to genomic data as well.

**Table 1.**
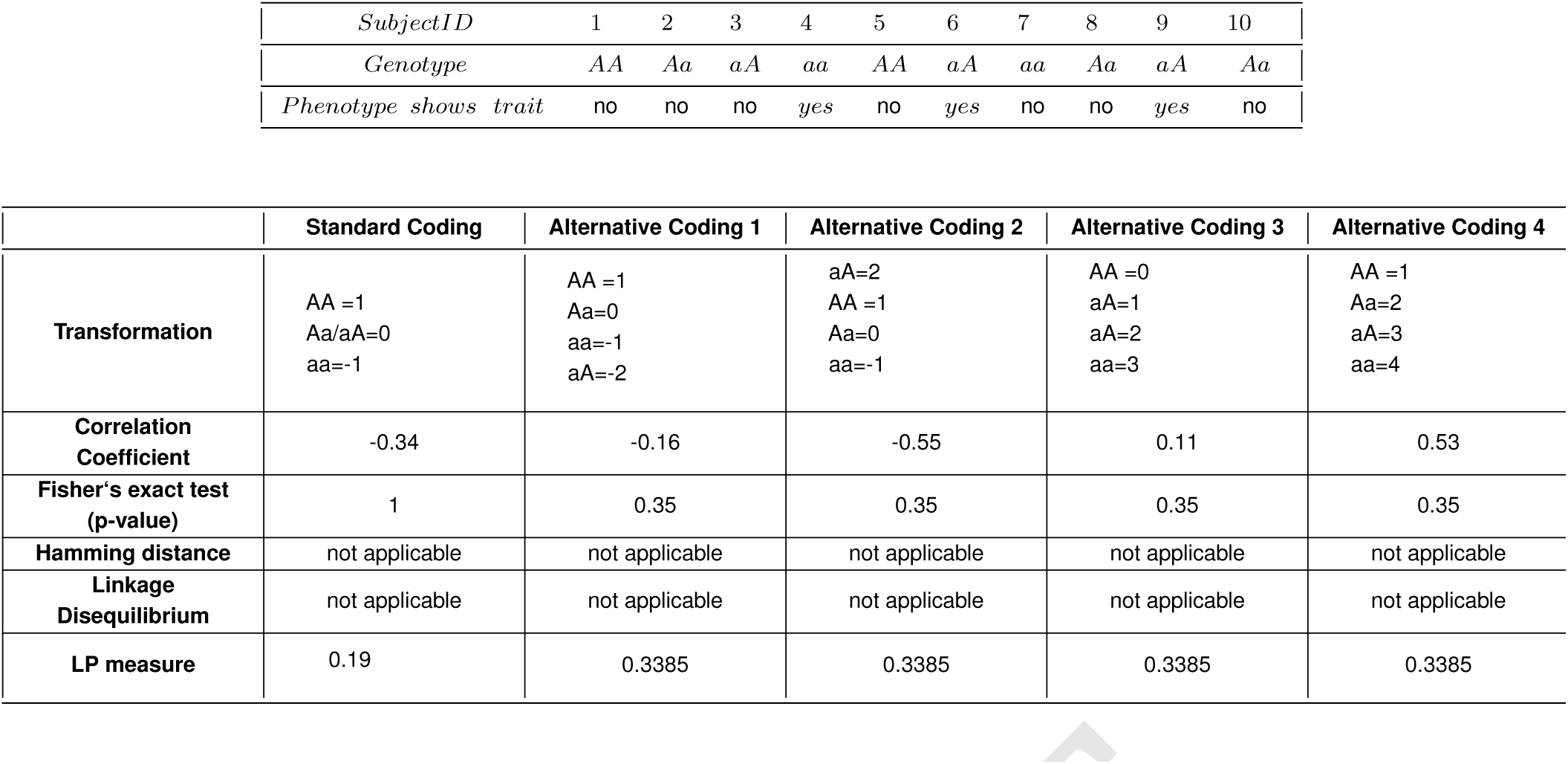
The upper table shows genotype-phenotype combinations for ten hypothetical subjects; the lower table provides several options for encoding the genotypic variants into numeric values (first column shows the standard coding) together with the respective correlation coefficients, p-values, LD distance, Hamming distance values and **LP** measure score.

**Errors introduced by the underlying model** are another issue that we will explore in greater depth using the example of INTERSNP(14, 18), a popular software tool used for GWAS of case-control SNP data and quantitative traits. The algorithm behind INTERSNP belongs to the class of linear Euclidean measures and models relationships between genomic variables such as *Y*, *X* and non-genomic variables *U* as a linear regression (when variables *U* are assumed to be explicitly available as data - i.e. U is *not latent*). However, linear regression is subject to a *model error* if the actual relationship between the variables is unknown and non-linear, or if some of the variables are latent or missing. This implies that even if all linear regression coefficients could be correctly determined from data that is available, the model may still lead to an incorrect and biased understanding of it.

**Finally, the concept of the p-value** that is integrated in many measures frequently leads to faulty results in GWAS and linkage studies. The main problem lies in the fact that p-values can only indicate the degree to which specific data are incompatible with a specific statistical model, but they do not measure the probability that the hypothesis in question is true, or that the given statistical model fits the data, even though these latter two questions are the very questions of interest (25).

All these problems arising with standard measures are additive and corrupt interpretations obtained from genomic data when, for example, trying to understand the differences between contributions of epigenetic/steric (19), populational and selection-driven (26) factors to LD.

As a means to addressing all of these problems, we propose an unbiased and computationally-tractable method called Linkage Probability (LP) to study the relationships between SNP by inferring information about all eventual relationships between the allele categories. This measure provides the posterior probability of a relationship between two categorical data sets while taking into account various forms of potential bias arising from the latent variables. We use an example to show that this method additionally permits the extraction of latent 3D steric effects from purely 1D SNP data. This allows steric, selection-driven and population-related patterns in SNP linkage relationships to be disentangled in a manner that improves upon the standard measures that are currently used in genomics.

We also demonstrate that this approach is helpful in validating associations between SNP when considered in relation to measures of structural/epigenetic features of the genome that are known to influence the expression of genes, a dimension that is not generally considered in the computation of standard measures such as linkage disequilibrium and cross-correlation. Finally, we raise important issues related to the unbiasedness of **LP** in relationship to the initial coding of categories and demonstrate that the use of this approach is entirely feasible in terms of the computational cost it entails.

## 1. RESULTS

### The **LP** measure

LP is the probability of (pairwise) relationships between SNP by inferring Bayesian relatoins between all categories of alleles, while taking into account the possibility that latent/unknown factors influence the outcome. **LP** is therefore based on an estimation of the posterior Bayesian relation probability between categorical datasets *Y*^*t*^ and *X*^*t*^, where the relations are measured in terms of the inferred conditional probabilities for *Y* and *X* and taking into account an eventual presence of latent effects *U*. Essentially, the suggested measure is computed in two steps:

(*Step 1*) First, the relations are quantified in terms of matrices (*m* × *n*) of conditional probabilities for available data *Y* and *X*; and

(*Step2*) Secondly, these matrices are used to obtain a single scalar posterior probability for the existence of a relation between *Y* and *X*.

An extended description of this procedure is given in the Materials and Methods section of this article. The entire methodology is explained in details in the Supplement.

Furthermore, the **LP** score takes into account unknown/latent variables and reveals a stronger relationship between the linkage of genetic variants and genomic contact strength compared to standard methods

#### Application to the analysis of human SNP data

Here, a practical application of the **LP** measure will be demonstrated using a dataset from the Human Genome Diversity Project (HGDP)(7, 27–29). This data has been used as a basis for a number of Genome Wide Association Studies (GWAS), in search of relationships between specific combinations of SNP and diseases (see, for example, (2, 3, 19)). Another use has been to establish chronological and evolutionary relationships between different human groups in the context of population genetics (7, 8). We used the dataset to generate results for pairwise relation analyses of SNP computed according to three methods: Cross-correlation measure (CC), the Linkage Disequilibrium measure (LD) and the Linkage Probability measure (LP). The first two measures are contingent on the implicit stationarity assumption (i.e., an assumption that the relation between SNP does not vary between individuals *t* = 1,…,*T*). To ensure that the comparison with the **LP** measure introduced here is fair, we deploy a variant of the **LP** measure computation using the same stationarity assumption (the details of computing the **LP** measure using the stationarity assumption are provided in section 3 of the Supplement). To provide a clearer analysis of the differences between the inferences made about patterns of relations using these methods, we considered a specific domain of chromosome 21 and calculated the SNP-SNP relation matrix for the Amyloid Precursor Protein (APP) gene, and its surrounding noncoding region (including 1500 kbp upstream and the same amount of sequence downstream of the APP sequence). Fig.1 illustrates the SNP-SNP relations inferred for the APP gene region using calculations based on the three methods CC, **LP** and LP. SNP relation matrices obtained for the entirety of Chromosome 21 is given in the Supplemental Figure. S1.

**Fig. 1.**
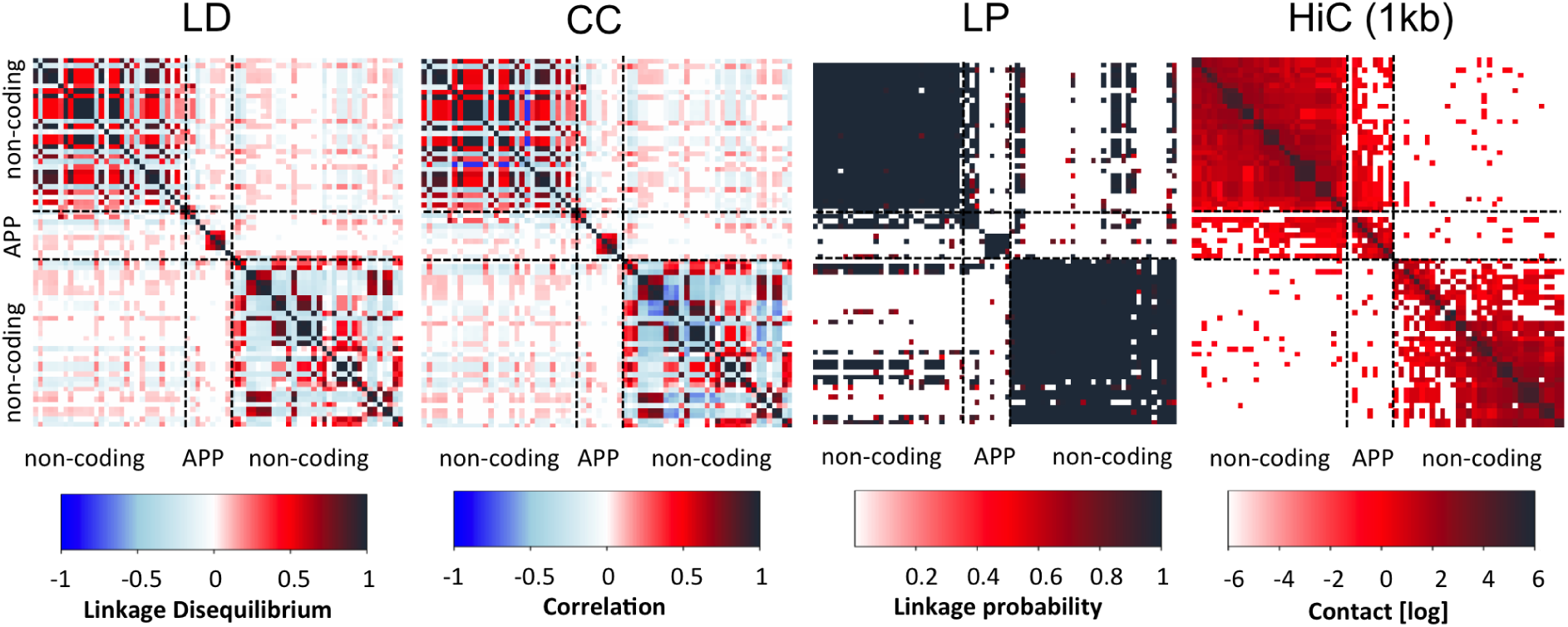
Pairwise relations for SNP located within 1500 kbp upstream and downstream of the the APP gene, calculated using (a) Linkage disequilibrium (LD), (b) cross-correlation (CC) and (c) Linkage Probability (LP) measures. Darkblue spots indicate a strongly negative relation, black spots indicate strongly positive relation and white spots null relations. Each black diagonal represents a calculation of the relation of a specific SNP with itself. Whereas the matrices for CC and LD are symmetric the matrix for **LP** provides the posterior probability of a relation between two SNP and is therefore not symmetric - despite the high similarity between the upper and lower regions of the chart. Subfigure (d) provides data from HiC at 1kb resolution for the same genomic region. The Spearman’s correlation coefficients for each pair are: (LP,HiC)=0.48, (CC,HiC)=0.07 and (LD, HiC)= 0.078, respectively.

The results obtained with CC and LD measures demonstrate very similar patterns of relation - with smaller blocks of related SNP aligned along the main diagonal and essentially no non-local relations apart from the diagonal, whether in the coding region nor in the non-coding area. The **LP** measure, in contrast, reveals a more complex picture of relations and at the same time provides values which are sharper than those obtained using the CC and the LD maps, because practically, it provides a close-to binary answer to whether or not an association is present. It shows crisp diagonal blocks of relations that are much larger – also in the coding region - and several non-local off-diagonal signals indicating a pairwise relation between SNP with a probability of almost one.

The two standard measures LD and CC yield contradictory results which consider only local SNP-SNP-relations, in comparison to results from LP, which estimate very strong probabilities for the existence of pairwise relations even for SNP that are separated by several hundred kbp, which is rather astonishing. In order to demonstrate that the patterns predicted with **LP** have biological meaning, we generated hypotheses that could be used to determine whether the distant linkages between pairwise SNP-SNP relations, exposed through the **LP** measure, could be associated with specific genomic features.

#### LP measure allows to reveal 3D steric effects in 1D SNP data

We computed the probabilities of relations between SNP as functions of a physical 1D-distance along the DNA sequence (measured in thousands of base pairs, kbp) for LD and LP. Since CC and LD produce highly similar results we selected LD and **LP** as a basis for the analysis. As can be seen from the Fig. 2, the function for the probability of a relation between two SNP as computed with the LD measure exhibits a monotonically decaying shape. The curve for low range interactions starts to drop very rapidly, descending further beyond distances of 10kb. As a result, most high linkage values are found between SNP separated by very short distances. This shape is in agreement with the generally accepted model that the probability of linkage between two SNP is very high for those in proximity to each other and decays extremely quickly and monotonously with an increase in the linear physical distance (> 50% reduction within a few hundred base pairs). This so-called LD decay rate is typically measured as the chromosomal distance (measured in base pairs) at which the average pairwise correlation coefficient (r2) drops to half its maximum value. (30–32). It is assumed that the shape of this curve reflects the high frequency of recombination events which create genotypic diversity in a population. The probability of contacts between SNP obtained through the **LP** measure, in contrast, reaches a plateau in the range of 1kb to 1Mb, followed by a drop and a plateau of a lower probability of contact over very long distances. We will demonstrate that this pattern is in better agreement with structural data than those obtained using LD and CC.

**Fig. 2.**
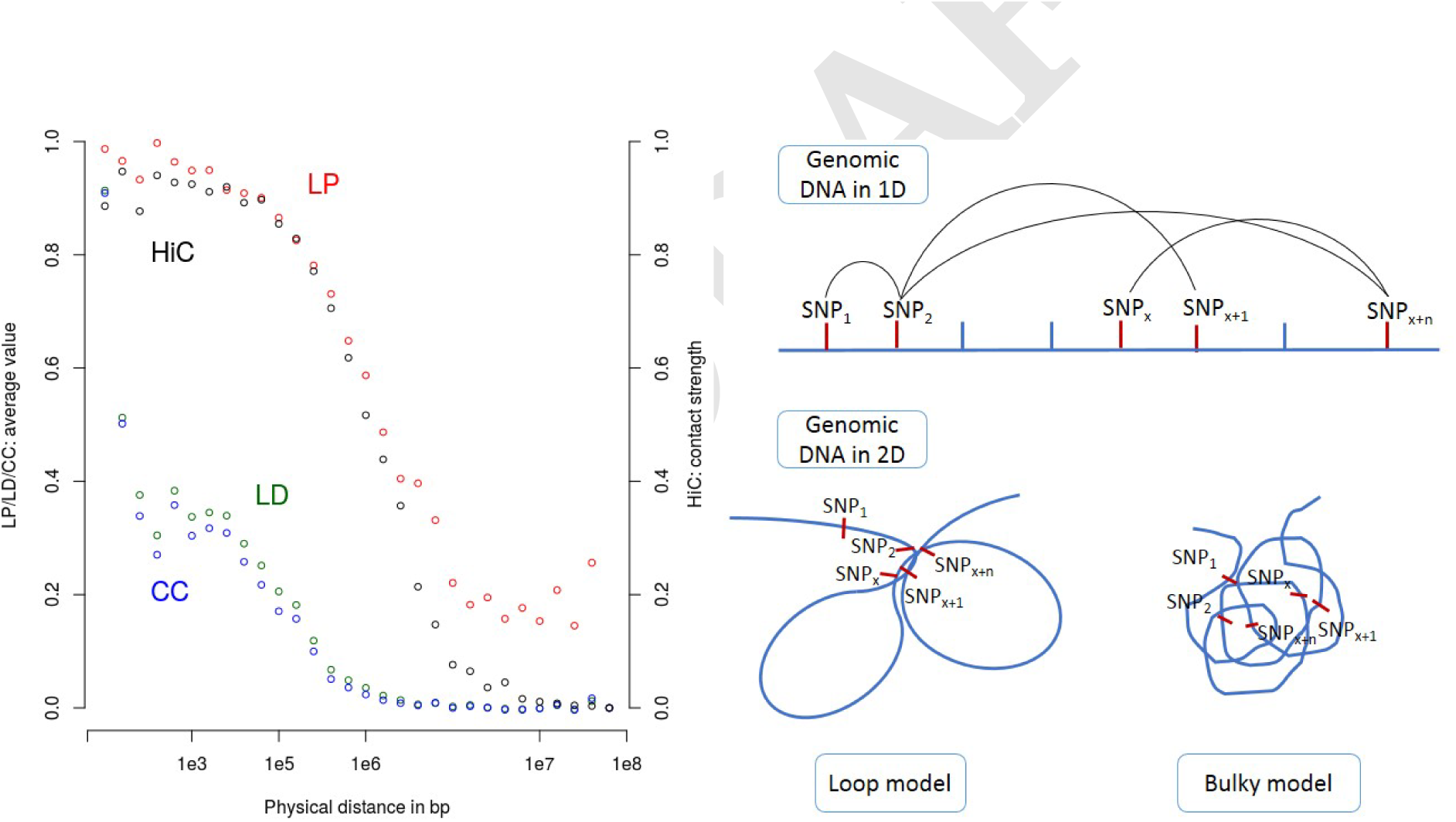
The relation between SNP linkage and epigenetic/steric features in 3D. Left: Average probabilities of relations obtained from **LP** (red), LD-measure (green) and cross-correlation(CC, in blue) in comparison to HiC values for the distances that separate two specific SNP (resolution of HiC data: 1kb). Right: Though SNP on the genome that are separated by fewer bps are spatially closer to each other, chromatin condensation and folding may alter this by bringing regions and SNP that are (linearly) farther apart into proximity. Two types of models can be used to describe chromatin domains: loop models (bottom left), which can favor linkage at intermediary distances, and bulky domain models (bottom right), which examines very compact structures and describes interactions at any short to intermediary distance. The initial plateau in the range 1e3-1e6 kb of distance for both **LP** and HiC scores is in favor of the bulky domain model. Plots built using SNP data from the 1000 genomes project showing similar trends can be found in the Supplemental Figure. S2

#### Genome in 3D

Several recent publications document not only the importance of the spatial and hierarchical organization of the chromosome architecture in the transcriptional control of genes (33–35) but also the existence of direct long-range interactions established between chromosomal sequences in so-called topologically associated domains (TAD) (36–38). This type of steric information can be obtained using genomic methods such as HiC (39), which probes the three-dimensional architecture of whole genomes and establishes 2D contact maps of genomic positions based on the sequencing of cross-linked sequences.

In order to verify our hypothesis that the observed maxima correspond to a DNA loop, we computed the distribution of contact intensity for the observed region using HiC data from (39). The HiC map for the APP region is shown in the very right panel of Fig1. It is evident that the **LP** measure is very close to matching HiC patterns (Spearman’s correlation coefficient=0.48) while both LD and CC maps were not able to detect any long distant SNP-SNP relations in regions with strong DNA-DNA contacts (red spots in the HiC map) at all. Both measures show a correlation coefficient of ≈ 0.07 with HiC data.

Figure 2A shows that both the intensity of contact (HiC value) and level of linkage (LP) are highly correlated and follow the same general shape, probably due to the fact that **LP** potentially succeeds at capturing latent factors such as genome structure. Interestingly, contrary to LD and crosscorrelation, which are very much in agreement, **LP** shows that the linkage is strong and similar for all SNP pairs in the low to intermediate distances, up to 1Mb, followed by rapid decay of linkage with distance. HiC follows the same pattern, which could mean that contrary to the classical loop model mostly proposed in the literature (40), our data can be fittet more exact by a model where domains are bulky, id est all genomic positions are close to each other in 3D space (Figure 2B). Further validations of these observed trends were found at least in short ranges (between 0 to 25 kbp) using SNP data from the 1000 genomes projects. For this, SNP data were sampled in define regions of 1Mb in order to make the computation of **LP** feasible in a reasonable amount of time. Similar trends were found in gene-rich and gene-poor regions (see Supplemental Figure. S2).

#### LP is not biased by initial coding of categories compared to cross-correlation and linkage disequilibrium

In the following, we shall provide a simple toy-example, aiming to reveal some important aspects of different relation measures relevant for more advanced genomic applications. Let us consider 10 individual subjects, either showing or not showing a certain symptom/trait. Concerning their genotype - for the purpose of simplification - we will consider differences in only one SNP position with four possible genotypic variants: homozygous major = AA, heterozygous = Aa or aA and homozygous minor = aa. The upper part of table 1 shows this data set of the 10 genotypes and phenotypes we are going to consider. The computational problem - mathematically expressed - can be worded as follows, along the lines of the terminology introduced above:

Two sets of data *X* = {*X*^1^,*X*^2^,…*X*^10^} and *Y* = {*Y*^1^,*Y*^2^,…*Y*^10^} (i.e., sample size *T* is 10) are given. For any subject *t* (*t* goes from 1 to 10), data *X*^*t*^ (genotype) takes values from *n* = 4 categories {*AA*, *Aa*, *aA*, *aa*}, and data *Y*^*t*^ (phenotype) takes data from *m* = 2 categories, {*yes*, *no*}. The question is to find if there is a relation between *X* and *Y*.

Values of relation measures obtained for this problem are summarized in table 1. The standard procedure of processing such categorical data for computational analysis starts by transforming (or coding) the genotypic variants {AA, Aa,aA, aa} as numbers { 1, 0, ‐1} or {0,1,2} (13, 14, 41). One assumption here is that *Aa* = *aA*, thus eliminating the need to discriminate between maternal and paternal information. In the following we shall start by using the first coding variant and calculate the correlation coefficient using the Matlab function **corr(X, Y)**. The correlation coefficient for the above example with standard coding is ‐0.34 ( implying an insignificant negative correlation). Another popular statistical test that can provide a measure of relation - the chi-square test (χ^2^) - is not applicable for this data set as it can only be performed if at least 80% of the cells have an expected frequency of 5 or greater, and no cell has an expected frequency smaller than 1 (42). These two conditions may be easily violated for genomics data as there may exist SNP that are very rare for certain phenotypes - resulting in very small values of the respective contingency table cells. Therefore, these two conditions of the standard χ^2^-test limit its applicability as a relation measure in large GWAS studies. Next, we chose Fisher’s exact test as a representative of the group of measures based on contingency tables. The two main advantages of Fisher’s test are: (i) it can be applied when the two conditions of the chi-square test are violated; and (ii) it provides an exact p-value - i.e., an exact conditional probability of observing this particular data (described by a contingency table with fixed sums of all rows and columns) if the two data sequences *X* and *Y* are independent (43).

To compute the p-values of the Fisher’s exact test, we used the Matlab functions from (43, 44), resulting in a p-value of 1 (the strongest possible outcome of the Fisher’s test, indicating that there is absolutely no relation at all between *X* and *Y*). LD and Hamming distances are not applicable to this example, as they can only be computed if *X* and *Y* are categorical variables, taking values from the same joint set of categories. For example, they can be straightforwardly used in the genomics context to which they apply for the analysis of relations between two different genotypes. They cannot be applied to infer the relations between the genotypes and the phenotypes. The lowest row of this table presents the results of the **LP** measure that is introduced above. The **LP** measure with a standard coding indicates that a relation between *X* and *Y* exists with a probability 0.19. At this point, it is important to stress the difference in the meaning of the standard p-values obtained with standard statistical tests (e.g., with a Fisher’s exact test) and the **LP** measure. p-value in standard tests is defined as *p* = ℙ[{*X*, *Y*}|*H*_0_] - i.e., it is true as a conditional probability to observe these (or more extreme) two data sequences *X* and *Y* under the assumption that the null-hypothesis *H*_0_ (i.e., the hypothesis that *X* and *Y* are completely independent in all categories). In contrast, as shown above, the **LP** measure is defined as

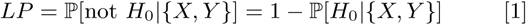

- i.e., it is the *posterior* probability of a null-hypothesis obtained after analyzing the data *X* and *Y*. Then, deploying the Bayes theorem we get:

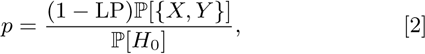

where ℙ[{*X*, *Y*}] is the *a priori* data probability (before looking at the data) and ℙ[*H*_0_] is the *a priori* probability of the relation model *H*_0_ (a model that is independent in all of the categories). A Step-by-step computation of the **LP** measure can be performed even manually for this data set. Please see the last section of the Supplement for a detailed step-by-step example.

#### LP is a transformation-invariant, user-independent, and computationally-tractable method to study linkage disequilibrium

Above we have considered different measures obtained with the standard transformation { 1, 0, ‐1} that uses the same coding 0 for the two mixed alleles {Aa,aA}. However, there may be scenarios where it would indeed make sense to discriminate between Aa and aA. However, it has been shown that certain epigenetic markers could be inherited over generations, leading to the involvement of epigenetic mechanisms in the intergenerational transmission of, e.g., stress effects or increased susceptibility to disease (41, 45–48). Therefore, integration of methylation-or epigenetic data into the GWAS-analysis would not only open completely new perspectives for such studies, but also require a more distinct discrimination between maternal and paternal chromosomes in the heterozygous genomic variants. Same holds true if there are more than three allele-variants. This happens very rarely but also demands the introduction of an additional category. This leads to a new problem: an infinite amount of possibilities for transforming/coding the four categories {*AA*, *Aa*, *aA*, *aa*} into numbers exists. They might be labeled as e.g. {1, 0,-1,-2} or {1,0,2,-1} or {1,2,3,4} or {0,1,2,3}, … and so forth. In the lower part of table 1 several transformation alternatives are given, together with the respective correlation measures, varying considerably from –0.55 (negative correlation) over 0.11 (no correlation) to 0.55 (indicating a positive correlation between the two datasets). All of these different values - implying completely different interpretations of a relation - are so different only due to the user-defined choice of coding in these four (out of an infinite number of other) variants. This means that the correlation coefficient is conceptually not appropriate for the analysis of relations between the intrinsically categorical data sets in genomics. In contrast, as can be seen from the table 1, both Fisher’s test, and the **LP** measure are invariant with respect to the user-defined choices of a transformation/coding.

Finally, we compare a computational cost of different measures for the artificial data sets of various lengths *T* (*T* is changing between 10 and 2’000) and with *n* = 4, *m* = 2. *Y*^*t*^ again contains phenotypes ‘yes’/’no’ and *X*^*t*^ the four genotypes, subject ID *t* is going from 1 to *T*. As can be seen from the Fig. 3, the computational cost is almost unchanged for the correlation measure and the **LP** measure. This observation is supported by a derivation shown in the Supplement, demonstrating that the computational cost of these two measures scales linearly in *T* . In contrast, the cost of relation measure based on Fisher’s exact test grows exponentially with *T*. This is due to the combinatorial explosion in the number of possible contingency tables that are needed to be generated, stored and processed (43, 44). Moreover, Fisher’s exact test assumes independence from latent factors, and would provide biased results when the impacts from latent factors (also known as confounding and population stratification effects (49, 50)) are not negligible.

**Fig. 3.**
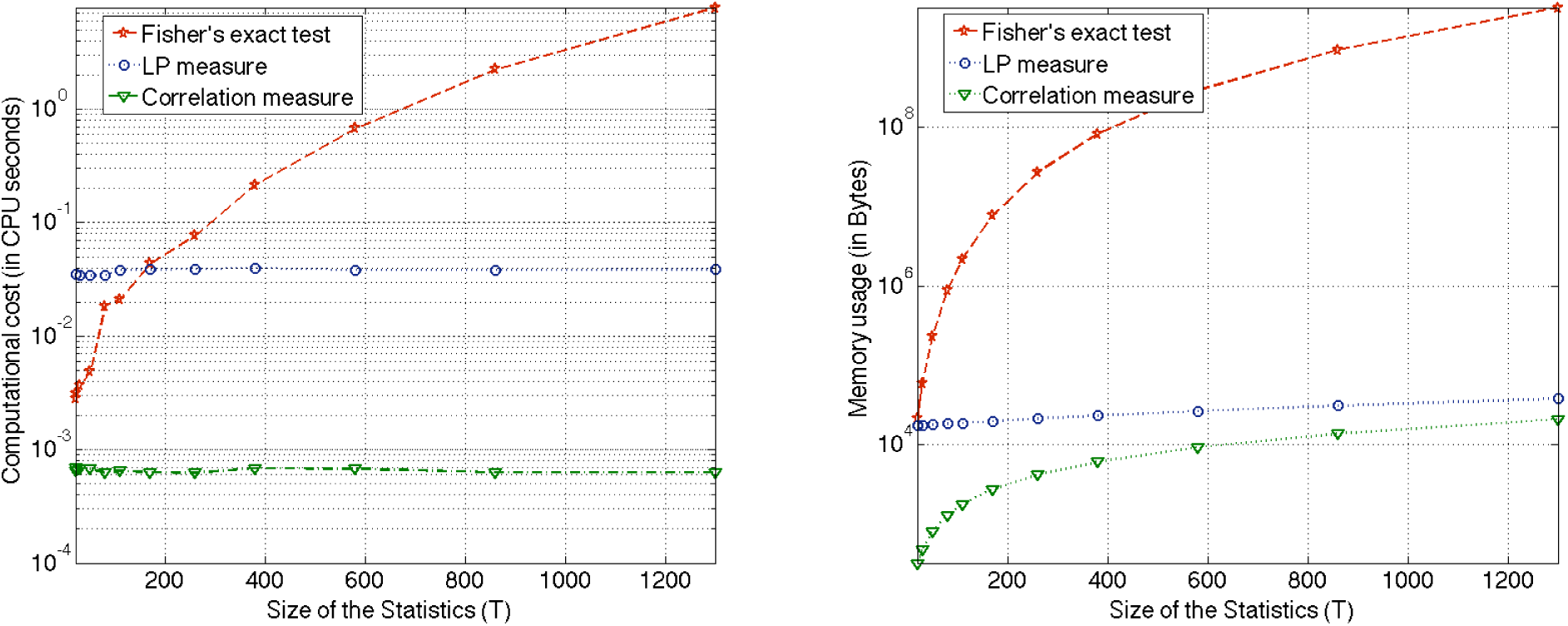
Computational comparison of different relation measures with respect to the computational time (left panel) and the memory usage (right panel). To assure the fairness of this comparison, all of the tests were performed on the same data sets, using the code implementation in the same programming language (Matlab) and the same PC (a MacBook Pro with a dual core processor).

It is straightforward to validate (please see the Supplement for more details) that the computational cost for the **LP** measure is scaling in the leading order as 𝒪(2^*m*^*mnT*), i.e., the **LP** relation measure is 2^*m*^ times more costly than standard measures such as covariance/correlation and linkage disequilibrium. This price, however, can be easily payed by parallelization. Results of this conceptual and computational comparison of different relation measures are summarized in the table 2.

**Table 2.** Summary: characteristics of the genomic linkage measure.

| | Transformation Invariance | Absence of a model error | Account for Latent Factors | Computational feasibility for large genomics applications ($n \geq 3, T \geq 1000$) |
| --- | --- | --- | --- | --- |
| Correlation | no | no | no | yes |
| Fisher's exact test | yes | yes | no | no |
| Information measures (e.g., Hamming-distance) | yes | yes | no | yes |
| LD measure | yes | no | no | yes |
| LP measure | yes | yes | yes | yes |

## 2. Discussion

We introduced a measure for linkage probability and illustrated its benefit - the possibility of obtaining a considerably more sharp and pronounced picture of pairwise SNP-relations (Fig. 1). Compared to other tools that are available for this purpose, the LP-measure - being based on the exact law of the total probability - is unbiased and designed to capture also putative latent variables as that may influence or bias GWAS studies. Furthermore, we compared different measures for the linkage of genetic variants with respect to aspects such as dependence on user-defined transformation/coding of the data, computational cost, and the possibility to take into account an eventual impact from latent factors and stratification effects (please see the tables 1, 2 and Fig. 3). We showed that the **LP** measure is less subject to model errors and explicitly takes into account an eventual implicit bias coming from the latent processes and/or stratification effects. Theoretical justifications for this measure were presented, including formal derivations, existence and uniqueness results, investigations of biasedness/unbiasedness in different parameter regimes, complexity estimates, and convergence properties (please see the Supplement). It was also shown (Fig. 3) that, despite of some additional computational effort needed for the inference of latent factors, this measure has the same leading order of computational complexity as the standard measures for linkage, thus making it potentially applicable to large realistic data sets in different areas of genomics.

The application of **LP** in comparison with Cross-correlation (CC) and Linkage disequilibrium (LD) to the phased HGDP data set from (7) demonstrated that **LP** is the only measure that is able to capture epigenetic/steric SNP association by revealing the latent proximity in 3D space (Fig. 2).

This means that the the **LP** measure has actually the potential to revolutionize GWAS studies since studies investigating also steric effects in GWAS have focused only on interactions between neighboring genes or enhancer-promoter loops (51–53). However, also TADs or chromatin hubs in which several regulatory regions are physically connected with their target genes can elicit a coordinated response(54–56). A disruption of such an entity through various means (e.g. via a point mutation) can cause significant domain rearrangements, inappropriate gene activation or human diseases (57). Disruptive SPNs in regulatory regions of such highly-coordinated elements may exert an dysregulative effect to several genes - thus leading to an impact that can be even more powerful than several mutation on the involved genes itself. Such a problematic disease-underlying elicitor in form of a SNP effecting the regulation of various genes, however could never be detected by normal GWAS or QTL-analysis, studis nor via the calculation of standard linkage disequilibrium or other pairwise analysis strategies. A systematic and unbiased integration of the chromatin structure and steric effect in GWAS, however, could be a major step towards understanding complex traits or disease.

## 3. MATERIALS AND METHODS

### Description of the **LP** measure

As a given datapoint, *X*^*t*^ can not simultaneously take two different values *x*(*j*_1_) and *x*(*j*_2_) (*j*_1_ = *j*_2_) at every given instant *t* (i.e., these events are *statistically-disjoint events*), relations between *X*^*t*^, *U*^*t*^ and *Y*^*t*^ can be expressed by means of the *law of the total probability*:

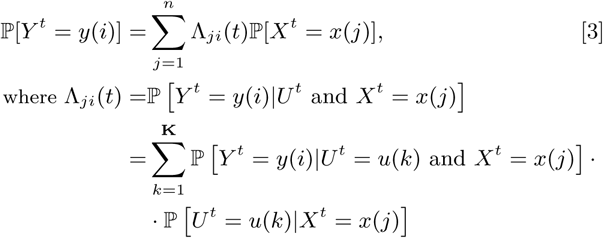

and where **K** refers to the a priori unknown number of categories (*u*(1),…, *w*(**K**)) of the latent impacts *U* (see section 1 of the Supplement for a detailed derivation). Expression [3] above is *exact* - in a sense that it is not subject to the modelling errors and holds for all linear as well as all non-linear systems.

In contrast to the above discussed linear Euclidean measures as implementend in INTERSNP, model [3] - being essentially a *law of the total probability* - is an exact model for all types of linear and non-linear relations between realizations *X*, *Y* and *U* when the coefficients Λ_*ji*_ are exact - i.e. it will provide an unbiased understanding of relations between the probabilities for *X* and *Y*.

If Λ_*ji*_(*t*) is available, it can also be used as an indicator for existence of causality relations between the processes *Y* and *X*: if {Λ}_*ij*_(*t*) ≡ ℙ[*Y*(*t*) = *y*(*i*)] for all *j* and all *t*, then the process *Y* is independent of *X* - meaning that information about process *X* provides no additional advantage in estimating the probability of the outcomes of *Y*. If {Λ}_*ij*_(*t*) ≠ ℙ[*Y*(*t*) = *y*(*i*)] - then there exists a causality relation between *X* and *Y* (58).

However, it is very straightforward to validate that the estimates of Λ obtained from the available data {*Y*^1^,…,*Y*^*T*^} and {*X*^1^,…,*X*^*T*^} by means of popular statistical tools, such as maximum log-likelihood approach, may be biased if the relation between the different categories *y*(*m*) of *Y* to *X* are not a priori known to be dependent or independent. For example, for *m* = 2 when Λ(*t*) is *t*-independent, if we know a priori that a certain category *y*(*m*) of *Y* is dependent on *X*, then the maximum log-likelihood estimator for Λ is 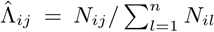 (where *N*_*ij*_ is the number of times *t* when *X*^*t*^ = *x*(*j*) and *Y*^*t*^ = *y*(*i*)). In the same situation, if we would a priori know that *Y* is independent of *X*, the maximum log-likelihood estimator for Λ will be different and take a value 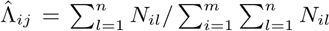. In general, different combinations of a priori dependence and independence relations between *Y* and *X* can be *encoded* by means of a Boolean vector *c*(*t*) = (*c*_1_(*t*), *c*_2_(*t*),…, *c*_*m*_(*t*)), where a particular vector component *c*_*i*_(*t*) takes the value 1 if the category *y*(*i*) is a priori assumed to be independent of *X* and the value *0* if a priori independence is assumed. It turns out that a particular maximum log-likelihood estimator of Λ(*t*) exists for each of these particular vectors^*^ *c(t*), computational algorithms for inference of the estimated relation matrices Λ̂(*t*) for different a priori choices of *c(t*) are described in sections 2 and 3 of the Supplement.

This procedure results in a set of *N* relation models (3) corresponding to *N* different combinations {*c*^(1)^(*t*),…, *c*^(*N*)^(*t*)} of a priori dependence and independence assumptions between *Y* and *X* (where the ordering of models is chosen such that *c*^(1)^(*t*) ≡ (0, 0,…,0) and *c*^(*N*)^(*t*) ≡ (1,1,…,1) for all *t* -i.e., ranging between the complete independence in all of the categories to a complete dependence in all of them). Yet, after performing this first step, we are facing the two following problems: (i) we obtain a set of *N* (*m* × *n*) matrices, but we need to get a single *number* that would characterize the overall relation between *X* and *Y*. (ii) We do not really know ahead what are the dependence and independence relations between the categories of *Y* and X and what is the "best" *c*(*t*) - as a matter of fact, this is what we actually want to find out.

To resolve both of these issues, we suggest adopting a Bayesian approach in a second step of the proposed measure computation. First of all, we observe that the obtained *N* models represent a full set of disjointed model combinations (i.e., no other model combinations are possible and all of the models are mutually-exclusive - meaning that any particular category of *Y* cannot be simultaneously dependent and independent of *X*). The least biased a priori assumption is to set all of the prior model probabilities ℙ_prior_ [*c*^(*i*)^(*t*)] (where *i* = 1,…, *N*) to equal 1/*N* – i.e., to assume that all of the relation combinations are equally probable a priori (i.e., before obtaining the data *X*^*t*^,*Y*^*t*^ for analysis)^†^. Then, using the Bayes’ theorem and the *N* obtained parameter estimates Λ̂, one can compute the *posterior* model probabilities {ℙ_post_ [*c*^(1)^(*t*),…, ℙ_post_[*c*^(*N*)^(*t*)]}. There are many possibilities to compute these posterior probabilities: in this manuscript, we chose a way to deploy a computation of the Bayesian Information Criterion (BIC) values (59, 61) for each of the *N* models (3), followed by a computation of the posterior BIC weights. Algorithmic details of the deployed procedure are explained in the section 4 of the Supplement.

Finally, after obtaining these posterior model probabilities, the measure of relation between *Y* and *X* can be computed straightforward as the posterior probability, namely that there exists at least some relation between these variables, i.e. as:

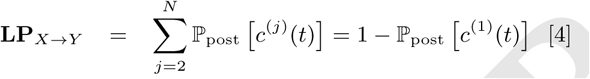

where *c*^(1)^(*t*) ≡ {0, 0,…,0}, corresponds to a model with not a single relation between the categories at any *t*. Thereby, **LP** is directly interpretable as a posterior probability of existence of at least some impact from *X* on *Y* in a presence of the latent/unobserved variables *U*. More specifically, 0 ≤ LP_*X*→*Y*_ ≤ 1.0 is taking the values close to 1, if there is strong evidence of a relation between the variables for a given data and being close to 0 if there is no such evidence from the data. Please also note that this measure does not rely on any external parameters and would provide results that are independent of any user-defined tuning. For further details and derivations of the **LP** measure please refer to the Supplement.

### Dataset used

For our analysis we used the phased HGDP data set from (7), containing 1272 SNP on the human chromosome 21 (regions 7 to 22) for 1107 individuals^‡^. Two particular data sets from this collection have been taken, representing a sample of (i) 149 Europeans and (ii) 149 individuals from sub-Saharan Africa. The ID numbers of selected individuals are available in the Supplement (file IID.csv). To compute the standard linkage disequilibrium measure *D*′ (normalized to the interval [‐1,1]) for these data sets, we have used the version 1.3.8.1 of the R-package ‘Genetics’ (62, 63). For computation of the cross-correlation measure, the data was first transformed deploying the standard encoding used in the literature - with +1 or 0 or ‐1 – denoting (respectively), whether an individual is homozygous with respect to the first allele or heterozygous or homozygous with respect to the second allele (64).

### Processing of HiC data

HiC data were downloaded from the NCBI webserver via ftp(http://www.ncbi.nlm.nih.gov/geo/query/acc.cgi?acc=GSE63525) published in (39). We chose to use the data generated from the immune cell line GM12878, because it is currently the only one available at 1kb of resolution and the authors have proven that the overall variation of genome structure is not very high. This meant that our conclusions would hold true for most of the somatic differentiated cell lines(file GSE63525_GM12878_combined_intrachromosomal_contact_matric Normalization was performed using a normalization vector generated by Rao et al. with the KR method.

### Computation of linkage disequilibrium and Spearman’s correlation coefficient

Linkage disequilibrium was computed using the R package *genetics version 1.7*, downloaded from the website of the CRAN. SNP were selected from the HGDP data set published in (7). SNP with low variability were removed from the analysis, leading to a set of 1226 SNP instead of 1272. Maps were generated using the default heatmap function of R. Spearman’s correlation coefficient was computed using the *cor(*) function from R (*"method=spearman"*). The data set is publicly available at the webpage of the N.Rosenberg’s Lab at Stanford: https://web.stanford.edu/group/rosenberglab/data

## ACKNOWLEDGMENTS.

The work of SG and DF was funded by the Center for Computational Sciences in Mainz (CSM), the work of IH was partially funded by the Mercator Fellowship in the DFG Collaborative Research Center 1114 "Scaling Cascades in Complex Systems" and by the Swiss Platform for Advanced Scientific Computing.

## Footnotes

* If vectors *c* are *t*-independent then there will be *N* ≤ 2^*m*^ models (3), if *c* is t-dependent then *N* ≤ 2^*mT*^ models - each characterized by a particular value of estimated matrix Λ. Section 3 of the Supplement presents a procedure to get a computationally-feasible approximation of the full subset of models *N* in a situation when *c* is *t*-dependent.

† According to the information theory, this uniform prior is the least biased prior - if no other a priori information is available (59). Moreover, due to the Bernstein-von Mieses theorem any particular choice of a prior becomes less and less important with the growing size *T* of the statistics - when all of the posteriors converge to the same posterior distribution (60). More background information for the relevant concepts from information theory is provided in the section 4 of the Supplement.

‡ This data set is publicly available at the webpage of the N.Rosenberg’s Lab at Stanford: https://web.stanford.edu/group/rosenberglab/data/huangEtAl2011/phased_HGDP+India+Africa

